# Perceptual integration without conscious access

**DOI:** 10.1101/060079

**Authors:** Johannes J. Fahrenfort, Jonathan van Leeuwen, Christian N.L. Olivers, Hinze Hogendoorn

## Abstract

The visual system has the remarkable ability to integrate fragmentary visual input into a perceptually organized collection of surfaces and objects, a process we refer to as perceptual integration. Despite a long tradition of perception research, it is not known whether access to consciousness is required to complete perceptual integration. To investigate this question, we manipulated access to consciousness using the attentional blink. We show that behaviorally, the attentional blink impairs perceptual decisions about the presence of integrated surface structure from fragmented input. However, when applying a multivariate classifier to electroencephalogram (EEG) data, the ability to decode the presence of integrated percepts remains intact when conscious access is impaired. In contrast, when disrupting consciousness through masking, decisions about integrated percepts and decoding of integrated percepts are impaired in concert with each other, while leaving feedforward representations intact. Together, these data show a dissociation between access to consciousness and perceptual integration.

**Significance statement:** Our brain constantly selects salient and/or goal-relevant objects from the visual environment, so that it can operate on neural representations of these objects. But what is the fate of objects that are not selected? Are these discarded, so that the brain only has an impoverished non-perceptual representation of them? Or does the brain construct perceptually rich representations, even when objects are not consciously accessed by our cognitive system? Here we answer that question by manipulating the information that enters into awareness, while simultaneously measuring cortical activity using EEG. We show that objects that do not enter consciousness can nevertheless have a neural signature that is indistinguishable from perceptually rich representations that occur for objects that do enter into conscious awareness.

## Introduction

Dating back to Helmholtz, conscious perception is thought to result from the unconscious integration of spatially scattered features, allowing the brain to make perceptual inferences about visual input (1). Historically, such mechanisms of integration have been linked to attentional selection (2). In this view, perceptual integration depends on conscious access, a position that echoes through in current theorizing about consciousness (3, 4). However, the link between perception and conscious access has been called into question in recent years, suggesting that perceptual structures may be formed despite not being consciously detected (5–7). In this counterview, conscious access does not play a causal role in perception itself, so that perceptual representations may exist without it.

The current study employs the Kanizsa illusion (see Fig. 1a), together with two well-known manipulations of consciousness, to assess whether neural representations can reach a state of integration in which features are combined to form perceptual entities, despite not being consciously detected. Kanizsa figures are similar to control figures in terms of physical input, but they have very different perceptual outcomes, notably an illusory surface region with accompanying contours (8) and increased brightness (9). These emergent properties are a primary demonstration of perceptual integration, as the constituent parts in isolation (the inducers) do not carry any of the effects that are brought about by their configuration.

**Figure 1.**
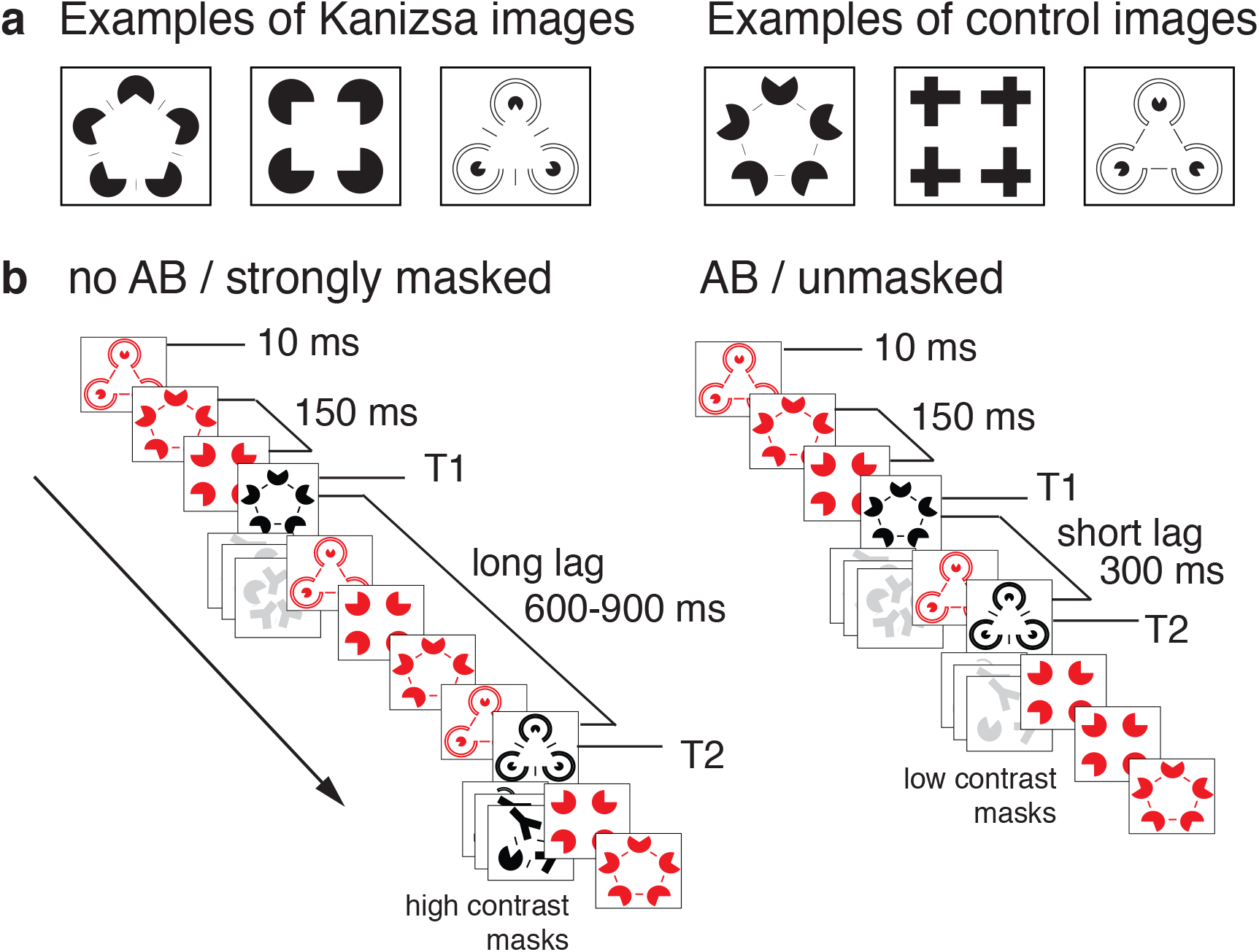
Experimental design. (a) Examples of different Kanizsa images and their controls as used in the experiment, see Fig. S1 for the complete stimulus set. (b) Examples of two of the four trial types in the factorial design: without an attentional blink (long lag) and strong masking (left) and with an attentional blink (short lag) and no masking (right).

Earlier work has shown that Kanizsa configurations can facilitate detection of target stimuli, with and without competing objects (10–14). However, in these studies conscious access has been implemented in various ways, while the dependent measure was always a behavioral response. The only study that has measured the neural substrate of perceptual integration in the absence of conscious report, postponed the behavioral response until after data collection (7), leaving open the possibility that subjects were consciously accessing the stimulus during scanning but had forgotten it at test time (15). The level at which conscious access and perceptual integration interact thus remains unclear. The current study employed several electroencephalographic (EEG) measures to investigate the neural substrate of perceptual integration under two very different types of manipulations known to affect consciousness: masking and the attentional blink (AB) (see Fig. 1b for the factorial design).

Masking is known to leave feedforward processing largely intact (16–20), while selectively interfering with perceptual integration and behavioral detection (14, 20–23) (see (24) and (25) for more in-depth reviews about the distinction between feedforward integration and recurrence-based perceptual integration). We therefore expected masking to simultaneously interfere with behavior and perceptual integration. The attentional blink on the other hand, is known to interfere with behavioral detection (26) and conscious access in particular (27, 28), but how it affects perceptual integration is not known. Therefore, the crucial question in the current experiment was whether perceptual integration would be impaired when access was disturbed by the attentional blink, as would be predicted when conscious access is causally involved in perceptual integration.

## Results

### T1 classification accuracy reflects perceptual integration

We recorded 64-channel EEG data from human subjects in two EEG sessions. Two black target figures (T1 and T2) were shown in a rapid serial visual presentation (RSVP) containing red distractors. Each target could either be a Kanizsa or a control figure (Fig. 1a and Fig. S1). T1 and T2 lag was varied, inducing an attentional blink at short lags (300 ms) with recovery at long lags (≥600 ms). In half of the trials, T2 was strongly masked using high contrast masks. In the other half, low contrast masks were used, so that there was no effect of masking (see examples of masks in Fig. S2). Examples of two of the four trial types are shown in Fig. 1b. At the end of each trial, subjects indicated whether T1 and/or T2 contained a surface region (see supplementary methods for details). The ability to distinguish surface from control figures was computed as the hit rate (HR) minus the false alarm rate (FAR), serving as a behavioral index of perceptual integration. T1 accuracy was high, at .90 (s.e.m. .02).

To establish a neural index for perceptual integration, we trained a linear discriminant classifier to categorize trials as either Kanizsa or control, using the amplitude of the EEG signal across electrodes as features for classification (see supplementary methods for details). To prevent response-related processes from contaminating our neural index of perceptual integration, the training set was obtained from and *independent* RSVP task containing Kanizsas and controls. In this task, subjects pressed a button on black figure repeats (1-back task on black targets while ignoring red distractors, see Fig. S3). This prevented response mechanisms from confounding classification performance, as target identity (repeat or not) was independent from stimulus class (Kanizsa or control), and all trials on which a response was given were excluded.

Next, we used the resulting Kanizsa classifier on the experimental runs, computing classification accuracy (HR-FAR, just as in the behavioral measure) for every time sample, yielding classification accuracy over time. As in behavior, Kanizsa versus control classification accuracy for T1 was well above chance, peaking at ~264 ms (Fig. 2a), and was strongly occipital in nature (see correlation/class-separability map (29) in Fig. 2b, supplementary methods for details). The fact that the classifier was able to discriminate Kanizsas from controls was reassuring, but we also wanted to establish a direct link between peak classification accuracy and perceptual integration.

**Fig 2.**
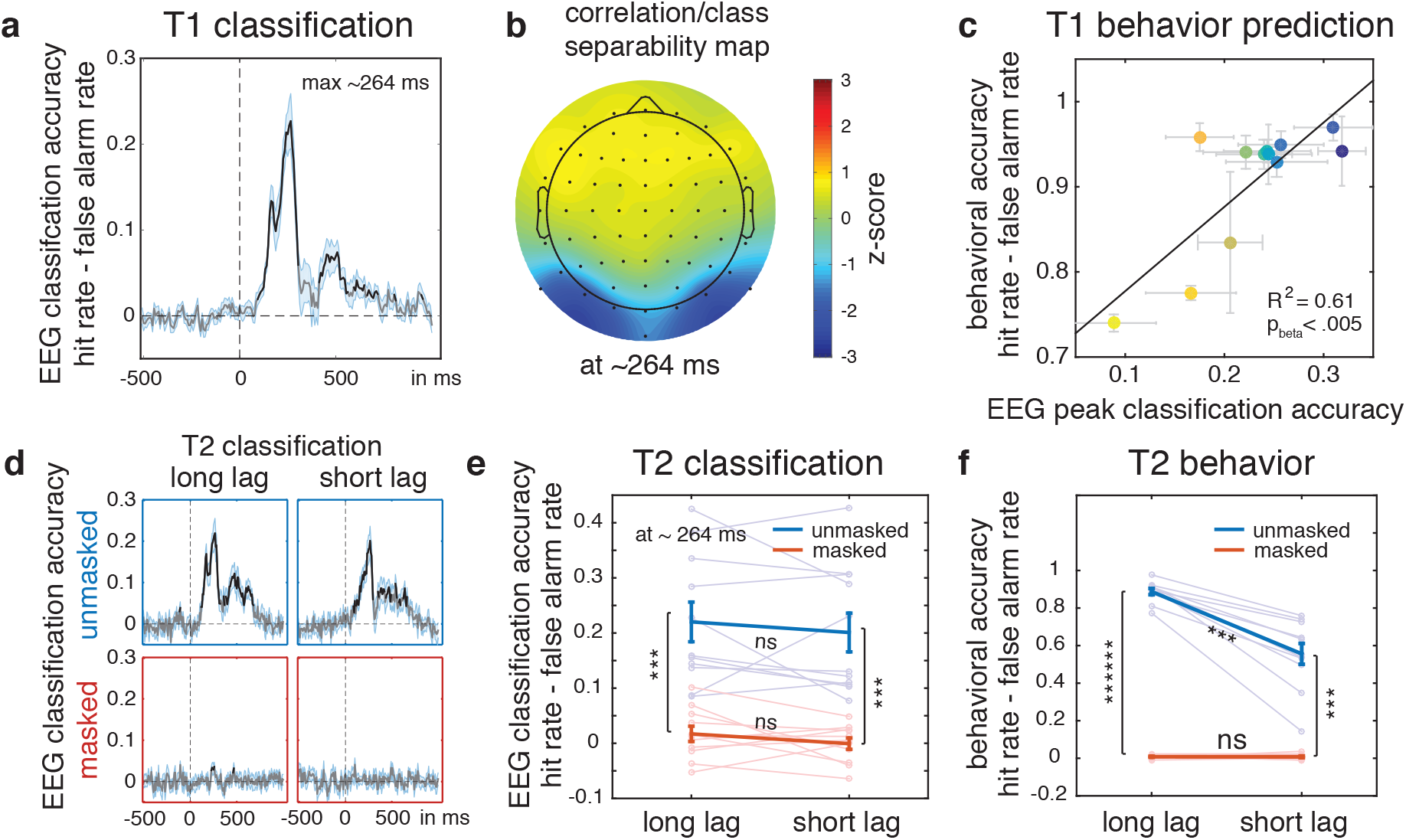
Peak classification accuracy reflects perceptual integration. (a) T1 EEG mean decoding accuracy of perceptual integration over time, black line reflects p<.05, +/− SEM in light blue. (b) and the correlation/class separability map reflecting the underlying neural sources for maximum decoding at ~264 ms, see methods. (c) The degree to which classification accuracy at ~264 ms predicts behavioral sensitivity to perceptual integration at T1 for the 12 Kanizsa-control pairs. Each colored data points is a Kanizsa-control pair, the color order follows the order of decoding accuracy in T1. For the full legend showing all pairs, see figure S1. (d) T2 EEG decoding accuracy over time for the four experimental conditions and (e) maximum decoding accuracy at ~264 ms for these conditions. (f) Behavioral sensitivity to perceptual integration for the four conditions (compare to e). Error bars are mean +/− SEM, individual data points are plotted using low contrast in the background.

To achieve this, we first computed behavioral accuracy separately for the 12 Kanizsa-control pairs that were used in the experiment. Different Kanizsa-control pairs yielded different behavioral accuracies due to inherent differences regarding the ease with which Kanizsa figures are perceptually integrated to result in surface perception (see Fig. S1 for the full Kanizsa-control stimulus set). Next, we applied a robust linear regression analysis (30) to determine whether T1 peak classification performance for these pairs (our neural index for perceptual integration) would be able to predict behavioral accuracy at T1. Peak classification accuracy was able to predict behavioral performance with remarkably high accuracy (R^2^ =.61, p<.005, see Fig. 2c, using colored dots to refer to the specific Kanizsa-control pairs in S1), providing independent evidence that peak classification accuracy captures the signal underlying perceptual integration. Moreover, we were able to do so using an independent RSVP training set that is not confounded by response or decision mechanisms (note that further down the manuscript we also perform analyses in which we train the classifier on T1 to investigate the impact of these mechanisms directly).

### The attentional blink and masking differentially impact behavioral and neural measures of perceptual integration

Next, we wanted to establish how the attentional blink and masking affect our neural marker of perceptual integration. In terms of behavior, we observed the classic deleterious effects of both masking (mask vs. no mask, *F*_1,10_=426.54, *P*<10^−8^) and the attentional blink (short vs. long lag, *F*_1,10_=51.89, *P*<10^−4^) on accuracy (Fig. 2f). There was also an interaction (*F*_1,10_=52.17, *P*<10^−4^), which was entirely driven by the difference between unmasked long- and short-lag trials (post-hoc t-test, *P* <10^−4^).

We hypothesized that if both masking and the attentional blink impact perceptual integration, they should both affect neural markers of perceptual integration in similar ways. To enable a direct comparison with behavior, we extracted classification accuracy in the four experimental T2 conditions. Fig. 2d shows the entire time course, Fig. 2e shows peak classification accuracy at 264 ms (latency taken from T1). A 2×2 analysis of variance (ANOVA) showed a highly significant main effect of masking (*F*_1,10_=37.68, *P*<.001), but no main AB effect (short vs long lag) (*F*_1,10_=2.16, *P*=.172), and no significant interaction between masking and AB (*F*_1,10_=0.02, *P*=.963). Post-hoc t-tests confirmed significant costs for masked versus unmasked stimuli for both long and short lag (both *P*<.001), but no significant differences between long lag and short lag (both *P*>.25).

Thus, while we observe a strong effect of masking in both brain and behavior, the classic AB effect only occurs in behavior. To further statistically underpin the differential effect of conscious access on behavioral and neural measures of perceptual integration, we entered both measurements into a large 2×2×2 ANOVA with factors measure (normalized behavioral / normalized neural), AB (yes/no) and masking (yes/no). The validity of treating neural and behavioral HR-FAR data as repeated measures of the same thing (i.e. classification of a perceptual object) is discussed in the supplementary methods section. In line with the other results, this analysis showed a three-way interaction effect driven by differences in behavioral and neural classification accuracies (*F*_1,10_=9.30, *P*=.012), as well as a two-way interaction between measure and AB (*F*_1,10_= 10.92, *P*=.008) but no interaction between measure and masking (*F*_1,10_=1.51, *P*=.247).

### Feedforward processing remains intact during masking

These data show that masking disrupts perceptual integration whereas the attentional blink does not. However, a concern might be that masking wiped out all processing of the stimulus, rather than specifically affecting perceptual integration, resulting in a floor effect. To test this, we selected a subset of the stimulus set that could be divided orthogonally according to its impact on input energy (contrast) or its impact on perceptual integration (surface perception). Fig. 3a illustrates this: the horizontal axis captures differences in perceptual integration (surface perception on the right but not on the left), while the vertical axis captures difference in bottom up energy (high contrast between the inducers and the background versus low contrast between inducers and background, see Fig. S5 and supplementary methods for a specification of the entire stimulus set). If masking wipes out all stimulus processing, we should no longer be able to classify high versus low contrast stimuli. We computed classification accuracy for feature contrast on the one hand and perceptual integration on the other, using a within-condition eight-fold cross validation scheme (see supplementary methods for details). The results are shown in Fig. 3b–c. In an early time window ~80-90ms, both masked and unmasked stimuli showed highly significant classification accuracies for feature contrast (left panes, masked: t(10)=7.45, *p*<10^−4^; unmasked: t(10)=8.82, *p*<10^−5^, statistics at ~92 ms, T1 peak latency). Thus, despite strong masking, the bottom-up signal is processed up to the point of contrast detection. Conversely, masking does wipe out classification accuracy on the perceptual integration dimension (right panes, masked: t(10)=−.19, *p*=.852; unmasked: t(10)=6.82, *p* <10^−4^). Note that for all analyses, the same type of masks would follow all stimulus classes (regardless of whether these were Kanizsa, control, high- or low contrast), such that the masks themselves could not bias classification accuracy. These results show that masking selectively abolishes perceptual integration, leaving feedforward processing largely intact, corroborating previous work (20, 21). In addition, this shows that the reduced classification accuracy for perceptual integration cannot be explained by a generic effect of reduced classification sensitivity under masking.

**Fig 3.**
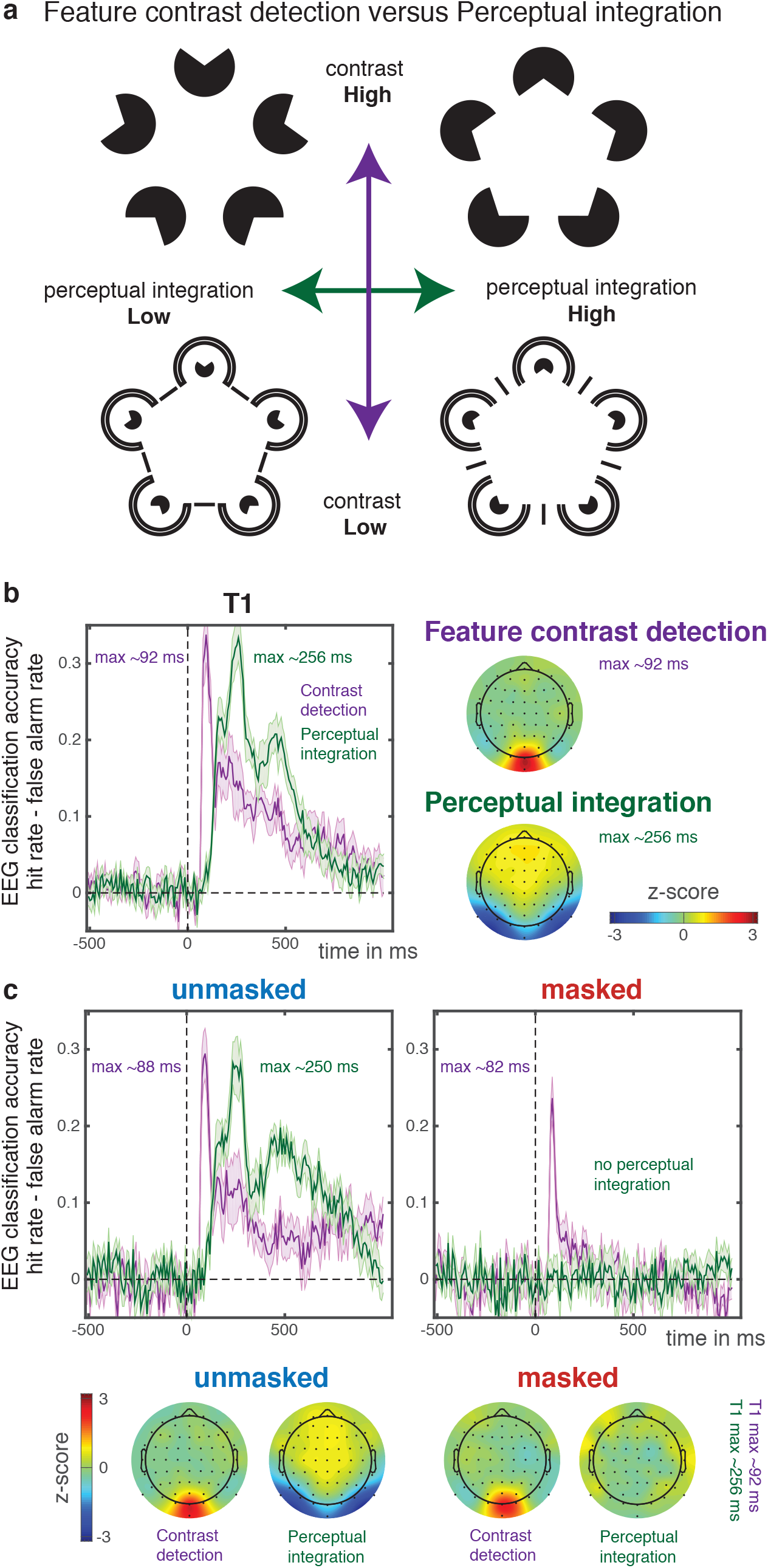
Separating out perceptual integration and feature contrast detection. (a) Example stimuli that were used to orthogonally classify feature contrast and perceptual integration on the same data. (b) Classification accuracies across time for contrast detection and perceptual integration (left) as well as correlation/class separability maps (right) for T1, (c) and for unmasked (left) and strongly masked trials (right). Line graphs contain mean +/− SEM.

### Masking selectively disrupts perceptual integration

Another concern might be that EEG classification accuracy is an all-or-none phenomenon whereas behavior relies on graded evidence. In such a scenario, the behavioral effects on perceptual integration (Fig. 2f) might not be reflected in classification accuracy (Fig. 2e) due to a lack of sensitivity of the classifier to smaller effects such as those observed during the AB. To test this hypothesis, we conducted a control experiment in which we used a staircase to titrate mask contrast to get a weaker behavioral effect of masking, similar in magnitude to the effect of the attentional blink in Fig. 2f (see supplementary methods for details). Fig. 4a shows the resulting behavioral effect of weak masking in this experiment. When computing classification accuracy on these data, we see that it nicely follows behavior (Fig. 4b-c), t(5)=3.82, *P*=.012. Together, these results show that the drops in behavioral accuracy caused by masking and the attentional blink have different root causes: masking impacts perceptual integration directly, whereas the attentional blink leaves it intact.

**Fig 4.**
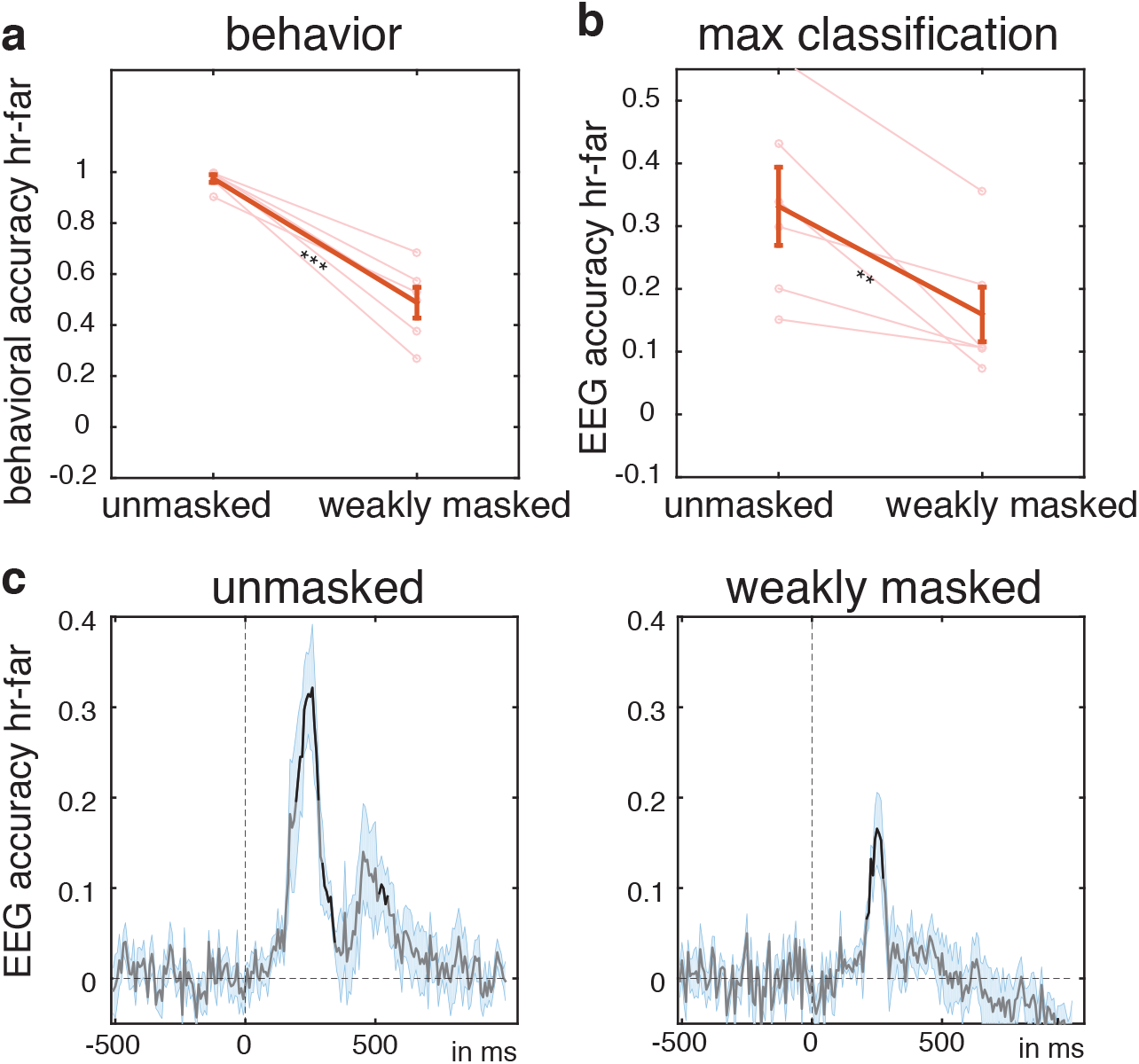
Masking control experiment. (a) behavioral results (b) max classification accuracy. Error bars are mean +/− SEM, individual data points are plotted in light in the background. (c) Raw decoding accuracies over time for unmasked and weakly masked conditions. Line graphs contain mean +/− SEM, black line reflects p<.05.

### Perceptual integration predates conscious access

So what neural process causes the dip in behavioral accuracy during the attentional blink? A natural hypothesis would be that the attentional blink interferes with conscious access after perceptual integration has already taken place. If true, we should be able to observe evidence of a selection process that results in conscious access at a later point in time. Investigating this issue requires a classifier that is sensitive to such a selection mechanism. Since the independent training runs that we used for training the classifier in the first analysis were designed to control for the direct influence of decision-related processes, these would not capture such a selection mechanism. The neural response to T1 however, does involve a conscious decision about the presence of a Kanizsa. We therefore trained a classifier on T1 data, and tested it on T2 data (see supplementary methods for details). Fig. 5a and 5b show classification accuracies for the four experimental conditions when using this T1 classifier.

**Figure 5.**
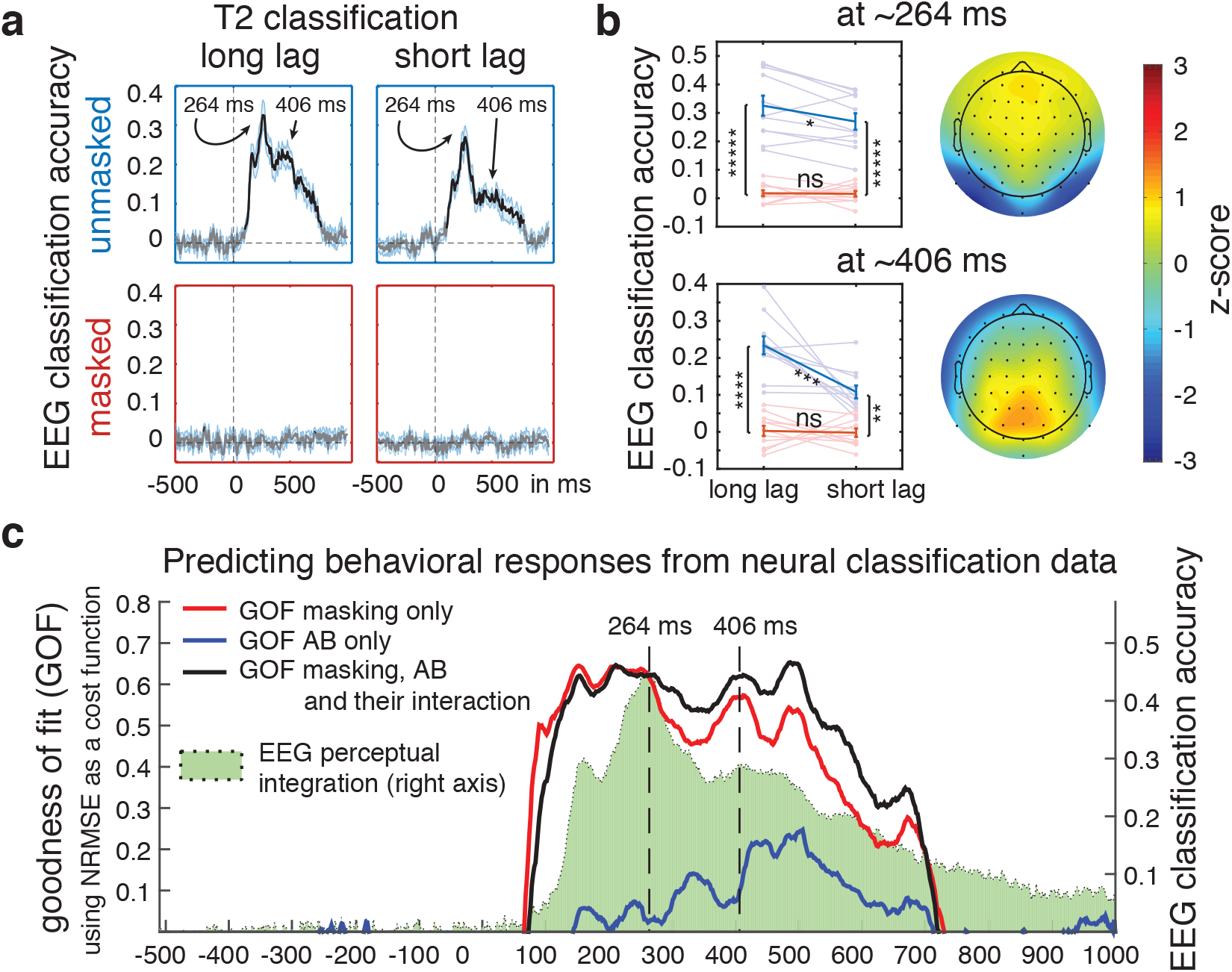
The impact of masking and AB on perceptual integration over time (a) EEG classification accuracy for the four experimental T2 conditions when training on T1. (b) EEG classification accuracies and correlation/class separability maps plotted at peak classification performance 264 ms (top) and at the second peak 406 ms (bottom). Blue lines represent the unmasked condition, red lines represent the masked condition. At the 264 ms time point, there was a strong main effect of masking (F_1,10_=91.63, P<10−5), a main effect of AB (F_1,10_=8.22, P=.017), and a trending interaction between masking and AB (F_1,10_=4.06, P=.071). To test directly whether the measurement source (neural or behavioral) at 264 ms results in a differential effect on classification accuracy, we entered the normalized measurements into a large 2×2×2 ANOVA with factors measure (behavioral/neural), AB (yes/no) and masking (yes/no), see online methods. There was no interaction between measure and masking (F_1,10_=.274, P=.61), but there was an interaction between measure and AB (F_1,10_=6.75, P=.027), as well as a trending three-way interaction (F_1,10_=4.50, P=.060), confirming that even when decision mechanisms are allowed contribute to classifier performance, the neural data at 264 ms cannot explain the pattern of results that is observed in behavior. The 406 ms time point on the other hand follows the same pattern as behavioral accuracy (see main text for statistics) and has a spatial distribution that is homologous to that of a classical P300. (c) An estimation of the goodness of fit when using the normalized EEG classification accuracy data as a model for the normalized behavioral detection data (left axis). Datasets are either collapsed over the AB dimension (GOF masking), over the masking dimension (GOF AB) or without collapsing over either dimension (GOF masking, AB and their interaction). T1 classification accuracy is plotted as a green shade in the background for reference (right axis). Not until after the perceptual integration signal has peaked at 264 ms does the black line overtake the red line, showing a postperceptual contribution of AB to behavioral accuracy.

We again find the initial peak at 264 ms that was described before. Despite the potential contribution of decision mechanisms to classification accuracy when training on T1, this peak follows a pattern that is similar to the pattern that we observed when training on the independent training runs (cf. Fig. 2e and Fig 5b, top), and which is not in line with behavioral accuracy (Fig. 2f, see caption of Fig. 5b for statistical tests). So at what point in time is the behavioral effect of the attentional blink reflected in the neural data? The most notable difference when training on T1, is a second peak in classification accuracy occurring around 406 ms, and which is heavily modulated by the AB (see top of Fig. 5a, supplementary methods and Fig. S7). At this time point, the pattern of results is identical to that obtained in behavior (cf. Fig. 2f and Fig. 5b, bottom). All manipulations had highly significant effects on classification accuracy: a main effect of AB (*F*_1,10_=7.96, *P=*.018), a main effect of masking (*F*_1,10_=130.19, *P*<10^−6^), as well as a strong interaction effect (*F*_1,10_=14.92, *P*=.003).

To again directly compare behavioral to neural data at 406 ms, we once more entered the normalized measurements into a large 2×2×2 ANOVA with factors measure (behavioral/neural), AB (yes/no) and masking (yes/no). The results show highly significant main effects of AB (*F*_1,10_=23.65, *P*<.001), masking (*F*_1,10_=528.18, *P*<10^−9^), as well as a strong interaction effect between AB and masking (*F*_1,10_=51.55, *P*<10^−4^), but importantly, now no two- or three-way interaction effects with measurement (neural/behavioral, all *F*_1,10_<3.08, all *P*>.110), underpinning the similarity between behavioral and neural data pattern at this time point. The correlation/class separability map at 406 ms (Fig 5b, bottom) has the same topology as that of a classical P300 (or P3b), which has been frequently associated with conscious access and perceptual decision-making (27, 31, 32). Our data provide converging evidence that neural signals around the time frame of the P300 reflect a post-perceptual signal that is involved in conscious access, rather than perceptual integration itself. What we unambiguously show is that perceptual integration precedes such conscious access.

In a statistical sense we have so far regarded behavioral and classification accuracy data as repeated measures of the same underlying perceptual object. Another approach would be to view neural mechanisms as the cause of behavioral outcomes, by assessing the degree to which the neural data are able to serve as a model for behavior across time. To do this, we used normalized classification accuracies as reference points to determine the goodness of fit (GOF) with normalized behavioral accuracies as test data. As a measure of goodness of fit, we used the normalized root mean square error (NRMSE) cost function given by:

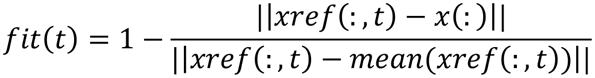

where *x* denotes the test data (behavioral accuracy), *xref* denotes the neural data (classification accuracy), || indicates the 2-norm (Euclidean length) of a vector, fit is a row vector of length *Nt* and *t* = 1,…,*Nt*, where *Nt* is the number of time points. NRMSE costs vary between -Infinity (bad fit) to 1 (perfect fit). If the GOF cost function is equal to zero, then *x* is no better than a straight line at matching *xref*. We obtained this fitness measure separately for the different factors by collapsing the neural and behavioral data either across the masking factor, across the AB factor, or without regard to either factor (see supplementary methods for details). The results are shown in Fig. 5c, where we also plot T1 classification accuracy as a reference for the time course of perceptual integration. Fig. 5c confirms that up to 264 ms, the masking manipulation uniquely models (predicts) behavior, indeed better than when AB is also allowed to contribute to the fit. Only after 264 ms does attention start to contribute to behavioral outcomes, trailing the perceptual integration signal itself and in line with prior analyses.

## Discussion

We show that EEG can be used to decode the presence of integrated percepts in visual cortex. Furthermore, we show that masking obliterates behavioral accuracy and classifier performance. Because the ability to decode feature contrast is retained under masking, the effects of masking on perceptual integration cannot be attributed to generic effects of masking on the sensitivity of the classifier. Rather, masking selectively disrupts perceptual integration while leaving feedforward signals intact (20, 21). Interestingly however, peak classification performance on integration remains unchanged during the attentional blink, despite causing a marked dip in behavioral accuracy. This shows that the brain is able to integrate features into perceptual objects when conscious access is impaired.

This conclusion is seemingly at odds with experiments on object-based attention. For example, in an experiment by Roelfsema and colleagues (33), monkeys were trained to perform a curve-tracing task in a display with overlapping curves. Attention to the task-relevant curve resulted in a spreading activation across V1 neurons that coded the features belonging to the curve, thus binding the constituent elements of the curve together. This suggests that serial access is the glue that unites an object, in line with the classical framework put forward by Treisman (2), and inconsistent with the position that conscious selection is not required for perceptual integration. Other studies have shown that such spreading activation follows Gestalt rules (34), and encapsulates task irrelevant features as long as they are part of a task relevant object (35, 36). However, with few exceptions, e.g. (37), task relevance and conscious access are intertwined in experiments on object-based attention. This suggests that the relationship between object-based attention and perceptual integration is caused by task relevance, rather than by conscious access per se.

Here we show that Kanizsa figures can be integrated in visual cortex despite not being promoted to a consciously accessible state. In contrast, masking destroys perceptual integration regardless of task demands. Naturally, this difference must be reflected in neural mechanisms. Dynamic feature grouping that underlies perceptual integration is thought to rely on cortico-cortical feedback (20, 24, 25, 38–44). While much remains to be learned about the origin of these feedback signals, evidence suggests that they originate from within visual cortex and are therefore local in nature (24, 39, 42–47). Although conscious access also involves feedback, this feedback originates from frontoparietal cortex (27, 48–53). In the consciousness literature, such long-range integration is often referred to as ‘global ignition’ (3, 54). The current data suggest that global ignition is not required to effectuate perceptual integration within visual cortex.

Our results also speak to a current debate about whether consciousness overflows cognitive access mechanisms (55, 56). In this debate, the question is whether access causes representational content to be extracted, or whether it acts to select from a rich representational set that cannot be accessed in its entirety, reminiscent of the debate on early versus late selection (57). In support of the latter position, a number of retro-cueing studies show that the representational capacity in early visual cortex is much larger than what can be accessed at any given moment, and that the extraction of this rich set from visual cortex does not require conscious access (58, 59). A recent study has questioned such results, suggesting that a retro-cue might serve to postdictively impact perception after the display has already disappeared, dismissing the idea that retro-cue experiments are able to convincingly show that perceptual representations can exist without access (60). The current experiment resolves this issue by using a direct neural measure of perceptual integration to show that perceptual integration precedes conscious access.

## Acknowledgements

We would like to thank Anouk van Loon for very helpful comments during the materialization of this project.

## References

1. Helmholtz Von H (1867) Handbuch der physiologischen Optik - Hermann von Helmholtz - Google Books.

2. Treisman AM, Gelade G (1980) A feature-integration theory of attention. Cognit Psychol 12(1):97–136.

3. Dehaene S, Changeux JP, Naccache L, Sackur J, Sergent C (2006) Conscious, preconscious, and subliminal processing: a testable taxonomy. Trends Cogn Sci 10(5):204–211.

4. Baars BJ (1988) A Cognitive Theory of Consciousness (Cambridge University Press).

5. Block N (2007) Consciousness, accessibility, and the mesh between psychology and neuroscience. Behav Brain Sci 30(5-6):481–99; discussion 499–548.

6. Lamme VAF (2010) How neuroscience will change our view on consciousness. Cogn Neurosci 1(3):204–220.

7. Vandenbroucke ARE, Fahrenfort JJ, Sligte IG, Lamme VAF (2014) Seeing without knowing: neural signatures of perceptual inference in the absence of report. J Cognitive Neurosci 26(5):955–969.

8. Heydt von der R, Peterhans E, Baumgartner G (1984) Illusory contours and cortical neuron responses. Science 224(4654):1260–1262.

9. Laeng B, Endestad T (2012) Bright illusions reduce the eye's pupil. P Natl Acad Sci USA 109(6):2162–2167.

10. Davis G, Driver J (1994) Parallel detection of Kanizsa subjective figures in the human visual system. Nature 371(6500):791–793.

11. Vuilleumier P, Landis T (1998) Illusory contours and spatial neglect. Neuroreport 9(11):2481–2484.

12. Mattingley JB, Davis G, Driver J (1997) Preattentive filling-in of visual surfaces in parietal extinction. Science 275(5300):671–674.

13. Conci M, et al. (2009) Preattentive surface and contour grouping in Kanizsa figures: evidence from parietal extinction. Neuropsychologia 47(3):726–732.

14. Harris JJ, Schwarzkopf DS, Song C, Bahrami B, Rees G (2011) Contextual illusions reveal the limit of unconscious visual processing. Psychol Sci 22(3):399–405.

15. Wolfe JM (1999) Inattentional Amnesia. Fleeting Memories, ed Coltheart V (MIT Press, Cambridge, MA), pp 71–94.

16. Di Lollo V, Enns JT, Rensink RA (2000) Competition for consciousness among visual events: The psychophysics of reentrant visual processes. J Exp Psychol Gen 129(4):481–507.

17. van Gaal S, Ridderinkhof KR, Fahrenfort JJ, Scholte HS, Lamme VAF (2008) Frontal cortex mediates unconsciously triggered inhibitory control. J Neurosci 28(32):8053–8062.

18. Rolls ET, Tovee MJ (1994) Processing speed in the cerebral-cortex and the neurophysiology of visual masking. P Roy Soc Lond B Bio 257(1348):9–15.

19. Kovacs G, Vogels R, Orban GA (1995) Cortical correlate of pattern backward-masking. P Natl Acad Sci USA 92(12):5587–5591.

20. Fahrenfort JJ, Scholte HS, Lamme VAF (2007) Masking disrupts reentrant processing in human visual cortex. J Cognitive Neurosci 19(9):1488–1497.

21. Lamme VAF, Zipser K, Spekreijse H (2002) Masking interrupts figure-ground signals in V1. J Cognitive Neurosci 14(7):1044–1053.

22. Wyatte D, Curran T, O'Reilly R (2012) The limits of feedforward vision: recurrent processing promotes robust object recognition when objects are degraded. J Cognitive Neurosci 24(11):2248–2261.

23. Koivisto M, Kastrati G, Revonsuo A (2013) Recurrent Processing Enhances Visual Awareness but Is Not Necessary for Fast Categorization of Natural Scenes. J Cognitive Neurosci 11:1–9.

24. Roelfsema PR (2006) Cortical algorithms for perceptual grouping. Annu Rev Neurosci 29:203–227.

25. Wyatte D, Jilk DJ, O'Reilly RC (2014) Early recurrent feedback facilitates visual object recognition under challenging conditions. Front Psychology 5:674.

26. Raymond JE, Shapiro KL, Arnell KM (1992) Temporary suppression of visual processing in an RSVP task: an attentional blink? J Exp Psychol Human 18(3):849–860.

27. Sergent C, Baillet S, Dehaene S (2005) Timing of the brain events underlying access to consciousness during the attentional blink. Nat Neurosci 8(10):1391–1400.

28. Vogel EK, Luck SJ, Shapiro KL (1998) Electrophysiological evidence for a postperceptual locus of suppression during the attentional blink. J Exp Psychol Human 24(6):1656–1674.

29. Haufe S, et al. (2014) On the interpretation of weight vectors of linear models in multivariate neuroimaging. Neuroimage 87:96–110.

30. Rousseeuw PJ, Leroy AM (2005) Robust Regression and Outlier Detection (John Wiley & Sons).

31. O'Connell RG, Dockree PM, Kelly SP (2012) A supramodal accumulation-to-bound signal that determines perceptual decisions in humans. Nat Neurosci 15(12):1729–1735.

32. Nieuwenhuis S, Aston-Jones G, Cohen JD (2005) Decision making, the p3, and the locus coeruleus-norepinephrine system. Psychol Bull 131(4):510–532.

33. Roelfsema PR, Lamme VAF, Spekreijse H (1998) Object-based attention in the primary visual cortex of the macaque monkey. Nature 395(6700):376–381.

34. Wannig A, Stanisor L, Roelfsema PR (2011) Automatic spread of attentional response modulation along Gestalt criteria in primary visual cortex. Nat Neurosci 14(10):1243–1244.

35. O'Craven KM, Downing PE, Kanwisher N (1999) fMRI evidence for objects as the units of attentional selection. Nature 401(6753):584–587.

36. Schoenfeld MA, et al. (2003) Dynamics of feature binding during object-selective attention. P Natl Acad Sci USA 100(20):11806–11811.

37. Peelen MV, Fei-Fei L, Kastner S (2009) Neural mechanisms of rapid natural scene categorization in human visual cortex. Nature 460(7251):94–U105.

38. Lamme VAF (1995) The neurophysiology of figure-ground segregation in primary visual cortex. The Journal of Neuroscience.

39. Fahrenfort JJ, et al. (2012) Neuronal integration in visual cortex elevates face category tuning to conscious face perception. P Natl Acad Sci USA 109(52):21504–21509.

40. Kok P, de Lange FP (2014) Shape perception simultaneously up- and downregulates neural activity in the primary visual cortex. Curr Biol 24(13):1531–1535.

41. Kok P, Bains LJ, van Mourik T, Norris DG, de Lange FP (2016) Selective Activation of the Deep Layers of the Human Primary Visual Cortex by Top-Down Feedback. Curr Biol 26(3):371–376.

42. Wokke ME, Vandenbroucke ARE, Scholte HS, Lamme VAF (2012) Confuse Your Illusion: Feedback to Early Visual Cortex Contributes to Perceptual Completion. Psychol Sci. doi:10.1177/0956797612449175.

43. Murray MM, et al. (2002) The spatiotemporal dynamics of illusory contour processing: combined high-density electrical mapping, source analysis, and functional magnetic resonance imaging. J Neurosci 22(12):5055–5073.

44. Halgren E, Mendola J, Chong CDR, Dale AM (2003) Cortical activation to illusory shapes as measured with magnetoencephalography. Neuroimage 18(4):1001–1009.

45. Hupe JM, et al. (1998) Cortical feedback improves discrimination between figure and background by V1, V2 and V3 neurons. Nature 394(6695):784–787.

46. Lamme VAF, Super H, Spekreijse H (1998) Feedforward, horizontal, and feedback processing in the visual cortex. Curr Opin Neurobiol 8(4):529–535.

47. Roelfsema PR, Lamme VAF, Spekreijse H (2000) The implementation of visual routines. Vision Res 40(10-12):1385–1411.

48. Corbetta MM, Shulman GLG (2002) Control of goal-directed and stimulus-driven attention in the brain. Nat Rev Neurosci 3(3):201–215.

49. Pooresmaeili A, Poort J, Roelfsema PR (2014) Simultaneous selection by object-based attention in visual and frontal cortex. P Natl Acad Sci USA. doi:10.1073/pnas.1316181111.

50. Gregoriou GG, Gotts SJ, Zhou H, Desimone R (2009) High-frequency, long-range coupling between prefrontal and visual cortex during attention. Science 324(5931):1207–1210.

51. Gregoriou GG, Gotts SJ, Zhou H, Desimone R (2009) Long-range neural coupling through synchronization with attention. Prog Brain Res 176:35–45.

52. Baldauf D, Desimone R (2014) Neural mechanisms of object-based attention. Science 344(6182):424–427.

53. Gross J, et al. (2004) Modulation of long-range neural synchrony reflects temporal limitations of visual attention in humans. P Natl Acad Sci USA 101(35):13050–13055.

54. Dehaene S, Sergent C, Changeux JP (2003) A neuronal network model linking subjective reports and objective physiological data during conscious perception. P Natl Acad Sci USA 100(14):8520–8525.

55. Block N (2011) Perceptual consciousness overflows cognitive access. Trends Cogn Sci 15(12):567–575.

56. Phillips I (2015) No watershed for overflow: Recent work on the richness of consciousness. Philosophical Psychology:1–14.

57. Deutsch JA, Deutsch D (1963) Attention - Some Theoretical Considerations. Psychol Rev 70(1):80–90.

58. Sligte IG, Vandenbroucke ARE, Scholte HS, Lamme VAF (2010) Detailed sensory memory, sloppy working memory. Front Psychology 1:175.

59. Vandenbroucke ARE, Sligte IG, Fahrenfort JJ, Ambroziak KB, Lamme VAF (2012) Non-Attended Representations are Perceptual Rather than Unconscious in Nature. Plos One 7(11):e50042.

60. Sergent C, et al. (2013) Cueing attention after the stimulus is gone can retrospectively trigger conscious perception. Curr Biol 23(2):150–155.

